# Single-particle measurements of filamentous influenza virions reveal damage induced by freezing

**DOI:** 10.1101/686642

**Authors:** Jack C. Hirst, Edward C. Hutchinson

## Abstract

Clinical isolates of influenza virus produce pleiomorphic virions, ranging from small spheres to elongated filaments. The filaments are seemingly adaptive in natural infections, but their basic functional properties are poorly understood and functional studies of filaments often report contradictory results. This may be due to artefactual damage from routine laboratory handling, an issue which has been noted several times without being explored in detail. To determine whether standard laboratory techniques could damage filaments, we used immunofluorescence microscopy to rapidly and reproducibly quantity and characterise the dimensions of filaments. Most of the techniques we tested had minimal impact on filaments, but freezing to -70°C, a standard storage step before carrying out functional studies on influenza viruses, severely reduced both their concentration and median length. We noted that damage from freezing is likely to have affected most of the functional studies of filaments performed to date, and to address this we show that it can be mitigated by using the cryoprotectant DMSO. We recommend that functional studies of filaments characterise virion populations prior to analysis to ensure reproducibility, and that they use unfrozen samples if possible and cryoprotectants if not. These basic measures will support the robust functional characterisations of filaments that are required to understand their roles in natural influenza virus infections.

## Introduction

While the virions produced by laboratory-adapted strains of influenza virus are commonly spherical, those produced by clinical isolates have varied morphologies (reviewed in (1) and (2)). They include spheres with diameters of 120 nm, bacilli with lengths of 200 nm, and filaments with lengths ranging to over 30,000 nm (1,3). Influenza viruses have been intensely studied due to their significant health impacts (4), but the role of filaments has been understudied and remains poorly understood (1).

At first glance, filament formation appears maladaptive; elongated filaments require more structural resources than spherical virions, and an equivalent mass of spheres would presumably be able to enter more cells. However, several studies suggest that filament formation is adaptive in natural influenza infections. First, clinical and veterinary isolates of influenza virus routinely form filaments when grown in cell culture. When these clinical strains are passaged in chicken eggs or cell culture they often lose the filament-forming phenotype, but when spherical laboratory strains are passaged in guinea pigs they gain it (5,6). Second, filament formation correlates with mutations conferring increased pathogenicity in the 2009 pandemic influenza virus, although it is challenging to separate filament formation from other effects on viral replication (7,8). Third, filament formation is common to several classes of enveloped respiratory viruses, including respiratory syncytial virus (9), parainfluenza virus type 2 (10), human metapneumovirus (11), and mumps virus (12), which suggests that filament formation is advantageous in the respiratory tract. Together, these findings suggest that filaments play a role in natural influenza infections that is dispensable or even maladaptive in cell culture. Identifying this role could reveal novel therapeutic strategies that would not be apparent from studying spherical laboratory strains alone.

Several suggestions have been made regarding the role of filaments. It has been suggested that filaments could traverse mucus better than spheres (13,14), that filaments could allow direct cell-cell spread in infection (15), or that filaments could initiate infections more robustly than spherical virions (16,17). None of these proposed roles have been shown to be clinically relevant. A major issue in identifying a role for filaments is that studies which focus specifically on filament properties often contradict one another. For example, some reports suggest that filaments are more infectious than spheres (16–18), while others suggest the opposite (3,19–21). Such discrepancies must be resolved if the function of filaments is to be understood.

It has been suggested that discrepancies between filament studies could arise from artefactual damage to the potentially fragile filaments during standard handling procedures (18). Concerns about such damage have been raised several times, with electron microscopy studies in particular often observing filaments that appeared to have been damaged from shear forces (3,7,16,18,20,22,23). However, this phenomenon has never been studied in detail and so uncertainty about the suitability of laboratory handling techniques persists. Characterising how filaments respond to routine handling is therefore necessary to remove this uncertainty and allow robust future investigations into their functional properties.

In this study, we aimed to determine whether common laboratory handling techniques could damage filaments. Using immunofluorescence microscopy and semi-automated image analysis, we measured the concentration and median lengths of high numbers of filaments and used these to assess the physical damage caused to filaments by a panel of common laboratory techniques. Most of the techniques we assessed did not cause substantial damage. A notable exception was the routine storage method of freezing, which significantly reduced the concentration and median length of filaments as well as inducing apparent capsid damage to the remaining virions. We show that the reduction in concentration and apparent capsid damage can be mitigated by freezing the samples in the presence of 10% DMSO, but the reduction in median length cannot. Together, our data suggest that most handling techniques are suitable for manipulating filaments but storing them using standard freezing procedures damages filaments and could skew functional assays into their properties.

## Materials and methods

### Viruses and cells

Mardin-Darby Canine Kidney cells (MDCKs) were cultured in Dulbecco’s Modified Eagle Medium (DMEM) (Gibco) supplemented with L-glutamine and 10% Fetal Calf Serum. Influenza A/Udorn/307/72(H3N2) virus (Udorn) was a kind gift from Prof David Bhella (MRC-University of Glasgow Centre for Virus Research) (3). To produce filament-containing stocks for analysis, confluent MDCK cells were infected at a multiplicity of infection of ∼1 and incubated in serum-free DMEM supplemented with 1 μg/ml TPCK-treated trypsin (Sigma) for 24 h. Supernatants were harvested and clarified at 1800 g at room temperature for five minutes unless otherwise specified.

Plaque assays were performed in MDCKs essentially as described by Gaush and Smith (24), with the agarose removed and cells stained with Coomassie blue to facilitate plaque counting.

### Virion manipulations

10 μl of Udorn-containing supernatant was added to 990 μl of PBS in a 1.5 ml microfuge tube before applying the mechanical manipulations of pipetting, vortexing, and sonicating. For pipetting, the entire sample was manually pipetted at 30 bpm using a Starlab 1000 μl pipette tip touching the bottom of the microfuge tube. Vortexing was performed at 2500 rpm using a Starlab Vortex. Sonication was performed at 50 Hz in a Kerry KC2 ultrasonic bath. For freeze-thawing, 1 ml of undiluted sample in a 1.5 ml microfuge tube (Greiner) was stored in a consistent position within a polypropylene cryobox (VWR), which was placed towards the centre of a C760 Innova – 70 °C freezer (New England Biolabs) for 1 h before being thawed in a 37 °C waterbath for approximately two min.

### Imaging

For confocal microscopy, samples were overlaid onto 1.3 cm coverslips, centrifuged at 1000 g at 4 °C for 30 minutes and fixed in 4% formaldehyde for 15 min before staining. Virions were labelled with the mouse anti-HA primary antibody Hc83x (a kind gift from Stephen Wharton, Francis Crick Institute) and goat anti-mouse Alexa-Fluor 555 secondary antibody (ThermoFisher). Coverslips were mounted using Prolong Diamond Antifade Mountant (ThermoFisher). 12 images from randomly selected sections of the coverslip were taken using the 63× oil immersion objective of a Zeiss 710 confocal microscope.

Image analysis was performed using FIJI (25). Images were auto-thresholded using the algorithm ImageJ Default. Particles with a circularity between 0.5 and 1 were removed using Particle Remover (26) to minimise the chances of cell debris being inaccurately scored as filaments. The number and lengths of the remaining particles were extracted using Ridge Detection (27,28). To assess particle distortion, the major axis and minor axes of the minimal fitted ellipse for each particle were calculated using Analyse Particles.

Estimated distributions of lengths within the population were calculated using a custom Python script. Graphs were plotted with ggplot2 (29,30) or matplotlib (31) and edited in Inkscape. All scripts are available at github.com/jackhirst/influenza_filament_analysis.

## Results

### Concentration and median length of filaments can be reproducibly measured by confocal microscopy

We aimed to assess damage to filaments by measuring the concentration and median length of filament populations before and after applying routine laboratory handling techniques. A procedure that entirely removed filaments should reduce the concentration, whereas a procedure that fragmented them should increase the concentration while reducing the median length. As assessing large numbers of filaments by electron microscopy is intensely laborious, we followed the example of previous studies which used immunofluorescence microscopy techniques to analyse influenza and RSV filaments (21,32,33). We centrifuged filaments on to untreated glass coverslips, taking advantage of the fact that filaments are known to adhere to glass (19). We could then visualise filaments by immunolabelling haemagglutinin (HA; Fig 1a). By applying a semi-automated ridge detection algorithm (Fig 1b), we were able to determine the length of filaments and the number of filaments in the micrograph, which we used as a proxy for their concentration.

**Figure 1:**
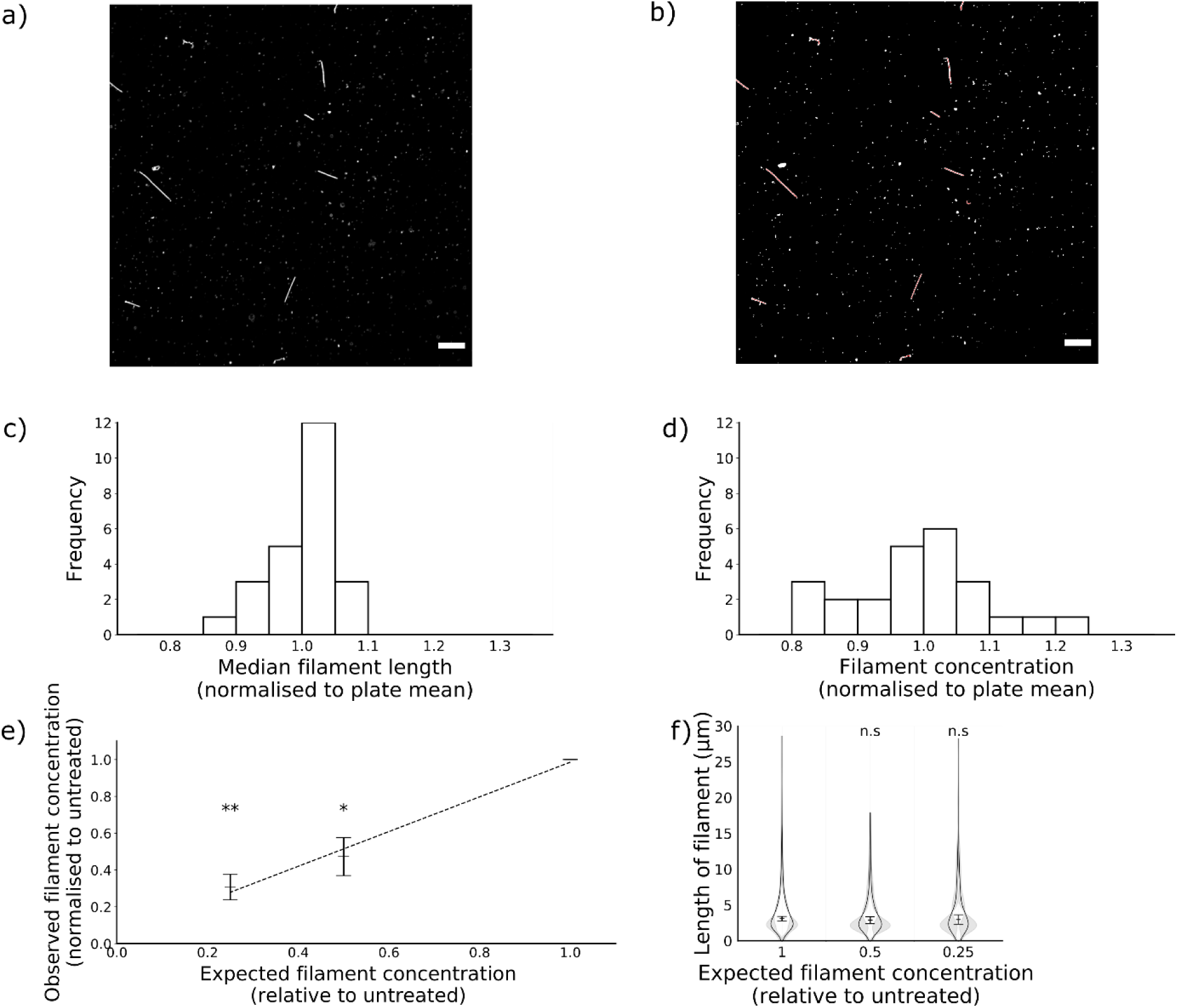
Concentration and median length of influenza virus filaments can be reproducibly measured by confocal microscopy. Influenza virus filaments were obtained by collecting supernatant from MDCK cells infected with the influenza virus A/Udorn/307/72(H3N2) for 24 h and centrifuging this onto glass coverslips. (a) To count and measure filaments, coverslips were immunostained for haemagglutinin and images were collected by confocal microscopy. (b) Next, a Ridge Detection algorithm was used to identify and measure filaments, indicated in red. (c,d) To assess reproducibility, three separate populations of filaments were divided into each well of 24-well plates. Measurements of length and concentration were taken from each well, and the mean for each of the 24 positions in the plate were calculated and then normalised to the total. Mean values for each position are shown of (c) median filament length within a well and (d) filament concentration per well. (e,f) To assess sensitivity, filaments were diluted in PBS prior to analysis. (e) Means and s.d. of filament concentration are shown of 3 experiments normalised to undiluted. Concentrations were compared to undiluted with two-tailed single-sample t-tests, * p <0.05, ** p < 0.01. (f) Frequency distributions of filament lengths were calculated for each sample. Violin plots indicate the mean frequency distribution, with the 95% CI shaded in grey. The median filament length was also calculated for each repeat; the means and s.d. of these median positions are indicated by lines and whiskers (n=3). Population medians were compared to the undiluted sample with two-tailed Student’s t-tests; n.s. = not significant (p>0.05).

To determine the reproducibility of the method, we assessed samples from the same stock of virus in every well of a 24-well plate. We calculated the average concentration and median length of each well in the plate across three repeats and normalised these values to the average of the whole plate. The concentration of filaments had a standard deviation of 0.1 as a proportion of the mean (Fig 1c) and the lengths of filaments had a standard deviation of 0.05 as a proportion of the mean (Fig 1d), confirming that this approach gave reproducible results.

To assess the sensitivity of the method, we altered the filament concentration by diluting samples in PBS. We found that the measured change in filament concentration matched the expected change (Fig 1e). Dilution should only affect concentration and not length, and indeed in both cases, the distribution of lengths in the filament population remained unchanged (Fig 1f). We concluded that immunofluorescence could detect changes in the concentration of a filament population over at least a four-fold range.

### Common laboratory manipulations do not substantially damage filaments

Having established a method to readily analyse the dimensions of filament populations, we could then compare the effects of various common laboratory manipulations on the stability of filaments. There are several plausible ways in which filaments could be destroyed or otherwise removed from a population. First, purification processes such as low-speed centrifugation to clarify virions from cell debris could inadvertently remove filaments. Second, mechanical manipulations of virions such as pipetting or vortexing could damage filaments through mechanical stresses or shear forces. Third, storing virions by freezing could cause damage due to changes in the chemical properties of the sample as it freezes or from ice crystals physically rupturing the membrane or capsid (34,35). In each of these cases, the elongated structure of filaments could make them more vulnerable than spheres and so more likely to be removed from the sample. We therefore tested routine handling techniques that could damage filaments in these ways.

First, we investigated clarification by low-speed centrifugation, which is commonly used to remove cell debris from virus samples. When we compared untreated samples and samples clarified at 1800 g for 5 min, we found no difference in filament concentration (Fig 2a) or median filament length (Fig 2b), suggesting that filaments were not being lost. To minimise the presence of HA-positive debris in our micrographs, all further experiments were performed using clarified samples.

**Figure 2:**
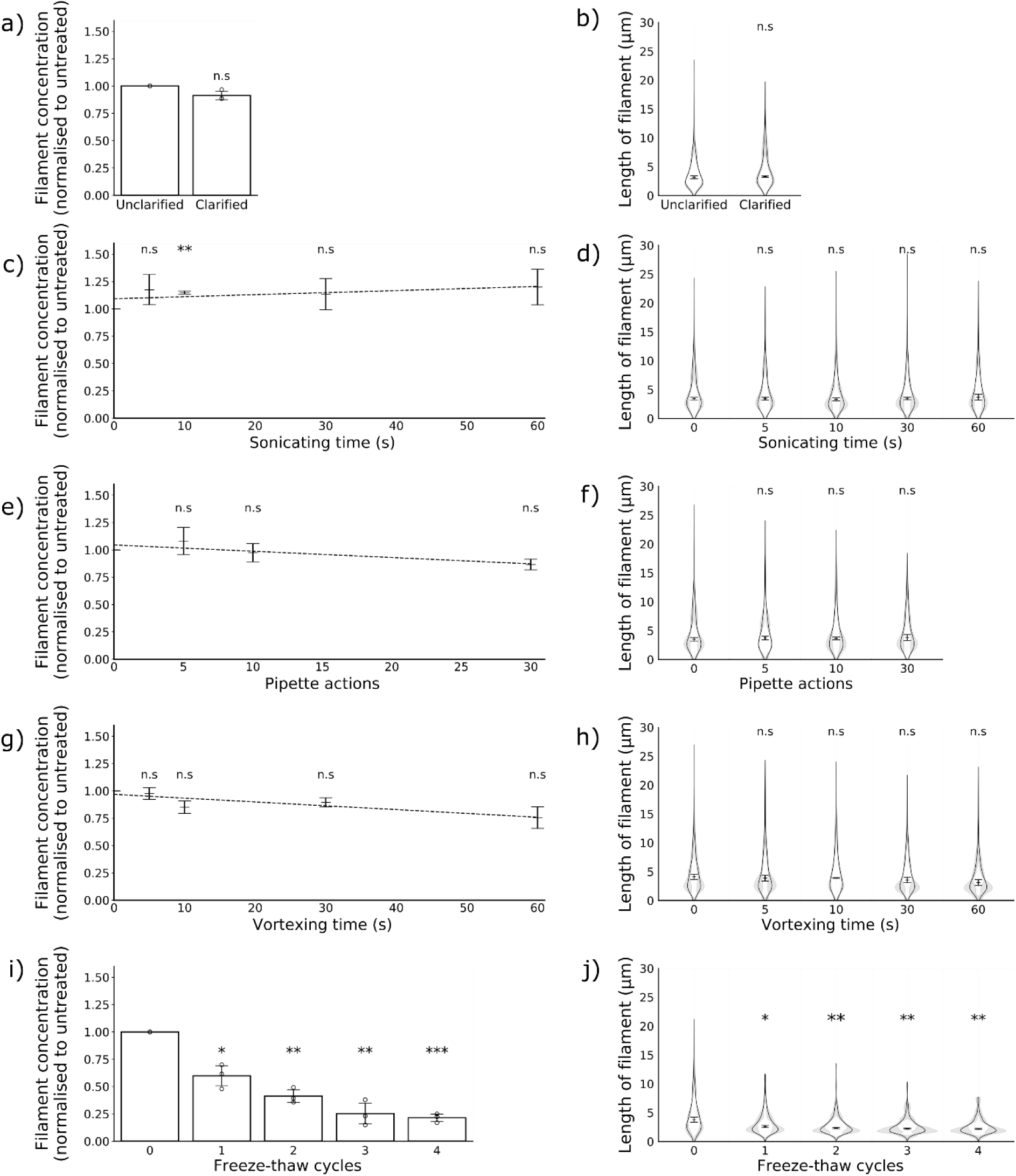
The effects of common laboratory manipulations on filaments. Concentration and length distributions of filaments in populations treated with clarification by low-speed centrifugation (a, b) and with increasing exposure to sonication (c,d), pipetting (e,f), vortexing (g,h) and freezing (i,j). (n = 3). Data from 3 repeats are shown. Concentration data are normalised to the untreated sample and the means and s.d. are shown; comparisons to untreated were made by two-tailed single-sample t-test: * p <0.05, ** p < 0.01, *** p < 0.001. Filament length distributions are shown as frequency distributions (mean, with 95% CI in grey) and distributions of the median filament length (mean position indicated as a line, s.d. as whiskers. Population medians were compared to the untreated sample by two-tailed Student’s t-tests: * p <0.05, ** p < 0.01, *** p < 0.001.

We then tested several common mechanical manipulations of virions: pipetting, vortexing and sonicating. We subjected samples to increasing levels of each treatment and compared the treated and untreated populations. We found that even after extended treatment, none of these techniques substantially altered the concentrations of filaments (Fig 2e, c, g) or the average filament length (Fig 2d, f, h). Together, these data suggest that mechanical manipulations do not cause substantial damage to filaments.

### Filaments are severely damaged by freezing

Finally, we investigated whether the routine storage method of freezing virus at – 70 to – 80 °C would damage filaments. We repeatedly placed virus either in a -70 °C ultrafreezer or a 4 °C fridge for one hour before thawing the frozen samples in a water bath at 37 °C for approximately 2 min and characterising the filament populations. We found that even a single freeze-thaw cycle reduced the concentration of filaments by almost half (Fig 3i) and median length of filaments by almost a third (Fig 3j). We observed further reductions in concentration and length with further freeze-thaw cycles. Freezing in this manner therefore causes severe damage to filaments.

**Figure 3:**
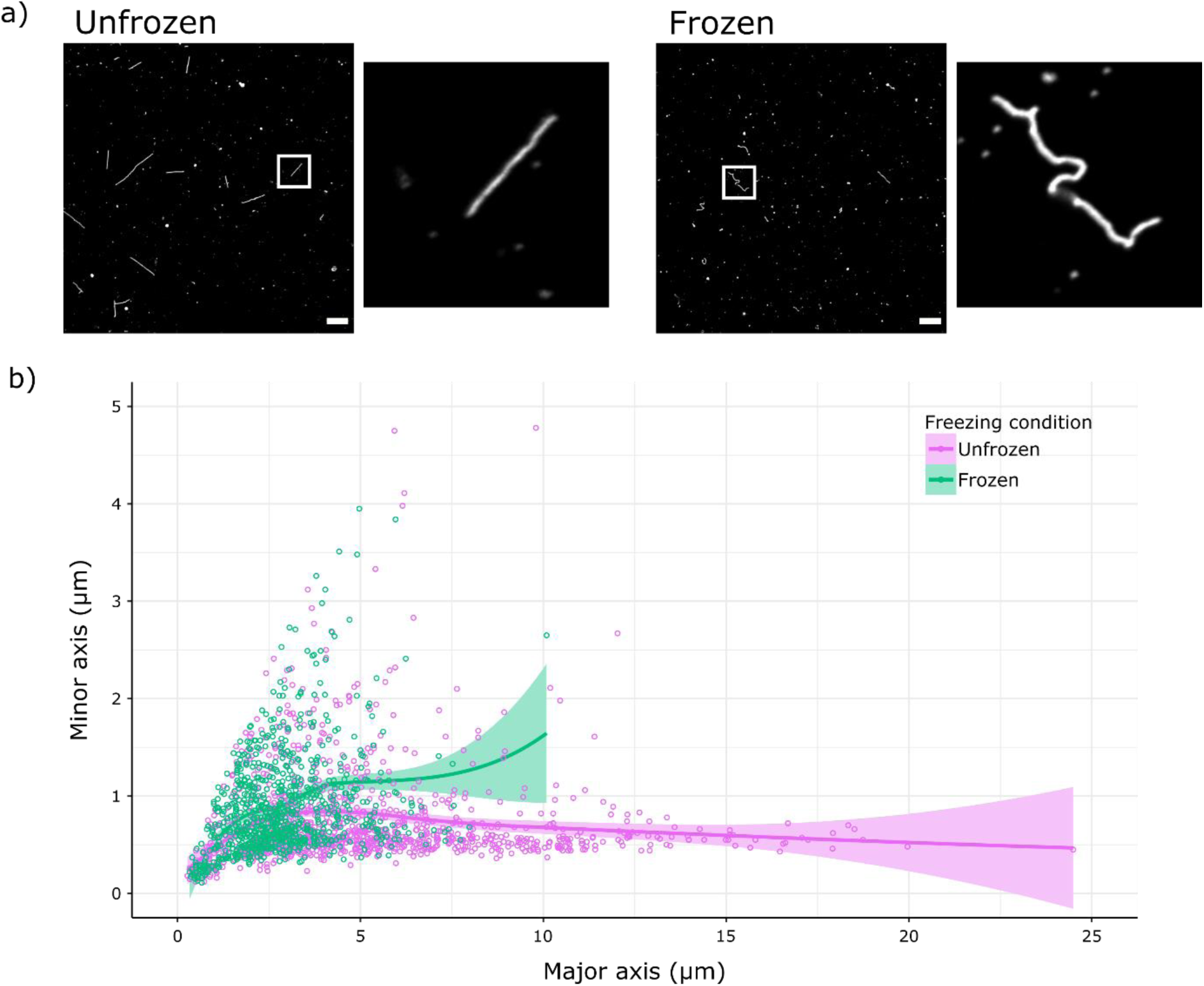
Freezing physically distorts filaments. (a) Representative images of unfrozen and freeze-thawed samples, immunostained for haemagglutinin and with insets magnifying an individual filament. Scale bar 10 µm. (b) Measurements of individual filaments from unfrozen samples and samples that had undergone a single freeze-thaw cycle, combining data from 3 separate experiments. Ellipses were fitted to each filament, and the major and minor axes of the ellipses are plotted, with the regression line (determined by local polynomial regression fitting) and 95% CI shown as a line and shaded area.

We also noted that the virions which had been frozen were often distorted along their length, suggesting damage to the viral capsid (Fig 3a). As the distortions compacted the filaments, we could quantify the distortion by fitting an ellipse to each filament and comparing the lengths of the major and minor axes. After a single freeze-thaw cycle the major to minor axis ratios were lower for frozen virions than unfrozen (Fig 3b). This suggests that even the virions that survived the freeze-thaw process were physically damaged.

Having shown that routine freezing could damage filaments, we sought an alternate freezing method that would minimise this damage. Snap freezing, and freezing in the presence of DMSO are commonly used to limit damage when freezing cells or tissue samples (36,37), so we reasoned that these might also reduce the damage incurred by filaments during freezing. When we compared these freezing methods with routine freezing, we found that both snap freezing and incorporating 10% DMSO prevented a reduction in filament concentration (Fig 4a). The major to minor axis ratios of fitted ellipses, indicating particle distortion, were reduced for snap freezing but not for freezing in the presence of DMSO (Fig 4b). However, the median filament length was reduced in all freezing conditions (Fig 4c). This suggests that while alternative freezing methods cannot entirely prevent freezing damage to filaments, incorporating DMSO can mitigate it.

**Figure 4:**
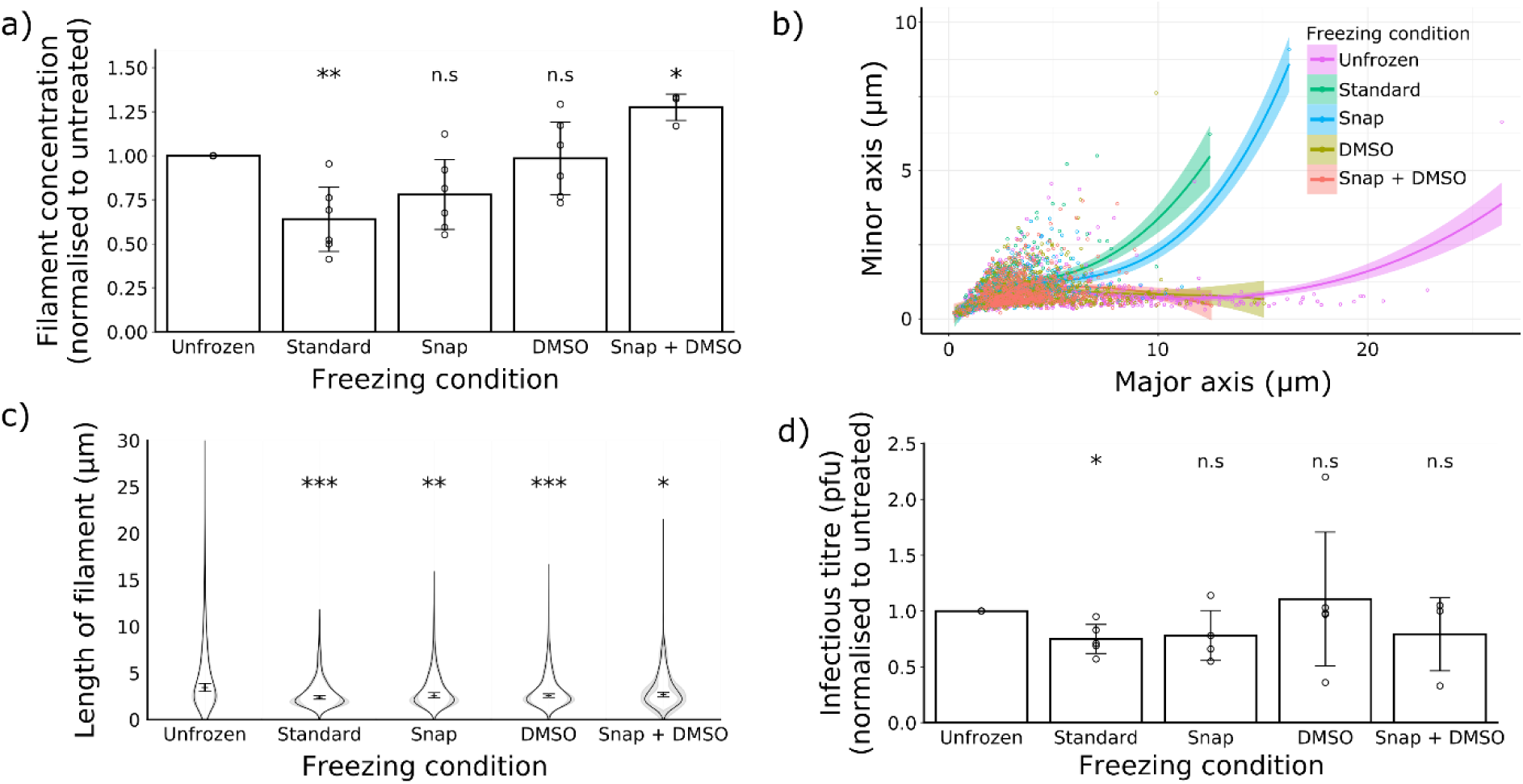
Alternative freezing methods can mitigate freezing damage. The effects of different freezing methods were compared for a single freeze-thaw cycle. (a) Filament concentrations after treatment, normalised to unfrozen. Means and s.d. of 3 repeats are shown, with comparisons to unfrozen by two-tailed one-sample t-test: * p < 0.05, ** p < 0.01. (b) Individual filament dimensions based on fitted ellipses, combining data from 3 separate experiments, showing regression lines (determined by local polynomial regression fitting) and 95% CIs as a line and shaded area. (c) Frequency distributions of filament lengths (mean, with the 95% CI shaded in grey) as well as the position of the median filament length (mean and s.d.). Population medians were compared to unfrozen with two-tailed, two-sample t-tests: * p <0.05, ** p < 0.01, *** p < 0.001 (n=6 except Snap + DMSO where n=3). (d) Infectious titres, measured by plaque assay in MDCK cells and normalised to unfrozen; means and s.d. are shown (unfrozen, standard, DMSO: n = 5; snap, n = 4; snap + DMSO, n = 3), with comparisons to unfrozen by two-tailed single-sample t-test: n.s. p > 0.05.

Finally, we considered whether the alternative freezing approaches could have different effects on the infectious titre of filament-containing samples, as freezing has been reported to reduce the infectious titre of influenza virus (34,38). Surprisingly, we found no substantial decline in virus plaque titre following any of the freezing conditions, including routine freezing (Fig 4d). Although unexpected, this result does suggest that the physical damage caused to filaments by freezing does not impact substantially on the infectivity of filament-containing stocks in tissue culture.

## Discussion

To determine whether common laboratory handling techniques could damage influenza virus filaments, we applied immunofluorescence microscopy to quantify the changes to filamentous virions caused by laboratory handling. We found that while clarification, sonication, pipetting and vortexing caused little or no damage, routine freezing substantially reduced the concentration and median length of filaments. We showed that the impact of freezing on filament concentration can be reduced by snap freezing or freezing in the presence of DMSO, but while DMSO could also prevent apparent capsid damage, no freezing method prevented the removal of long filaments.

Our data show that immunofluorescence microscopy can be used to assess changes to filamentous virion populations. Historically, determining filament numbers and dimensions has been attempted by manually counting particles using dark field microscopy (39,40), negative stain electron microscopy (17,41), or cryo-electron microscopy (3). The specific labelling of viral proteins in immunofluorescence microscopy makes it easier to automate virion detection, and thereby allows faster characterisation of larger samples than previous methods. Immunofluorescence microscopy also allows analysis of unconcentrated samples, which is challenging to accomplish with electron microscopy (14); furthermore it avoids damage or clumping that could be introduced by concentration procedures. The ability to rapidly assess the concentration of filaments in a stock also provides a major technical advantage when studying filament properties. Concentrations of filaments can vary between stocks, and studies of filament properties have not typically controlled for this.

The impact of freezing-induced damage on filaments could have been enough to skew previous investigations into their properties. Freezing is routinely used to store influenza virus samples (38), and previous studies on isolated filaments have often used frozen virions (18,41) or not explicitly stated their storage conditions (7,14,21,23,42,43). When using frozen samples, our data suggest the filament concentration could be almost half that of unfrozen, potentially reducing their contribution to a sample’s properties to below the limit of detection. The apparent capsid damage we observed also suggests that the surviving filaments may have different properties to their unfrozen counterparts. The possibility of freezing damage affecting results should therefore be considered when interpreting the current, contradictory literature of filament properties.

Based on our data, we recommend that future studies of influenza filament properties should avoid using frozen virus samples where possible. Using freshly prepared virus samples in assays should avoid the damage associated with standard freezing. If freezing cannot be avoided, freezing with 10% DMSO should reduce the damage, and microscopy can be used to assess the extent of any damage that has occurred. Avoiding artefactual damage in this way will make functional characterisation of filaments more robust, and so provide a firmer foundation for evaluating the role of filaments in infection.

As well as the influenza viruses, filament formation is common to several classes of enveloped respiratory viruses and our approach would be readily applicable to the study of these. The artefactual damage we observed with influenza filaments could affect these other viruses and so similar stability studies would also be relevant when designing functional assays for these viruses.

Although damage to filaments can cause problems when studying their properties, it may offer practical advantages in other contexts. Filaments can cause difficulties during influenza vaccine purification, as they can interfere with the filtration used to clarify allantoic fluid from infected chicken eggs (44). Simple treatments that remove or compact filaments in unpurified vaccine material could limit these difficulties. Freeze-thaw cycles could be an appealing approach for this, though for live-attenuated vaccines any advantages would need to be balanced against a potential reduction in infectious titre.

In conclusion, here we demonstrate a method for rapidly measuring and quantifying filamentous influenza viruses in unconcentrated stocks. This has intrinsic value in calibrating measures of filament properties, and by applying it to common laboratory manipulations we have shown that freezing can damage influenza virus filaments. We have also shown that snap freezing or adding a cryoprotectant can reduce freezing damage, but not eliminate it. This damage could explain discrepancies between past studies into filament properties. Our findings therefore remove a source of uncertainty present in filament research and provide a foundation for robust functional analyses of filaments in future. Such analyses will be necessary to finally identify the role of the clinically relevant but poorly understand filamentous influenza virions.

## Conflicts of interest

The authors declare that there are no conflicts of interest.

## Acknowledgments

We thank Amy Burke for assistance with microscopy. JCH is funded by an MRC QQR Core award to the University of Glasgow [172630], ECH is funded by an MRC Career Development Award [MR/N008618/1].

